# DirectContacts2: A network of direct physical protein interactions derived from high-throughput mass spectrometry experiments

**DOI:** 10.1101/2025.07.17.665435

**Authors:** Erin R Claussen, Miles D Woodcock-Girard, Samantha N Fischer, Kevin Drew

**Affiliations:** Department of Biological Sciences, University of Illinois at Chicago, Chicago, IL 60607

## Abstract

Cellular function is driven by the activity proteins in stable complexes. Protein complex assembly depends on the direct physical association of component proteins. Advances in macromolecular structure prediction with tools like AlphaFold and RoseTTAFold have greatly improved our ability to model these interactions *in silico,* but an all-by-all analysis of the human proteome’s ~200M possible pairs remains computationally intractable. A comprehensive cellular map of direct protein interactions will therefore be an invaluable resource to direct screening efforts. Here, we present *DirectContacts2*, a machine learning model that distinguishes direct from indirect protein interactions using features derived from over 25,000 mass spectrometry experiments. Applied to ~26 million human protein pairs, our model outperforms previous resources in identifying direct physical interactions and enriches for accurate structural models including ~2,500 new AlphaFold3 models. Our framework enables structural modeling of disease-relevant complexes (e.g. orofacial digital syndrome (OFDS) complex) offering insights into the molecular consequences of pathogenic mutations (OFD1) and broadly, establishes a highly accurate protein wiring diagram of the cell.

## Introduction

The self-assembly of proteins into protein complexes is critical for their biological function^1^. Disruptions in these protein complexes often give rise to human diseases including cancer^2^, developmental disorders^3^, and protein folding disorders such as Parkinsons^4^. Knowledge of each direct physical interaction of protein subunits (Figure 1A) ultimately gives rise to the wiring diagram of the cell. This diagram is essential for understanding the molecular mechanism of protein complex function including why protein complex function fails in pathological states. It also aids in drug target discovery^5^ and constructing three dimensional models of protein complexes on the proteome-wide scale.

**Figure 1:**
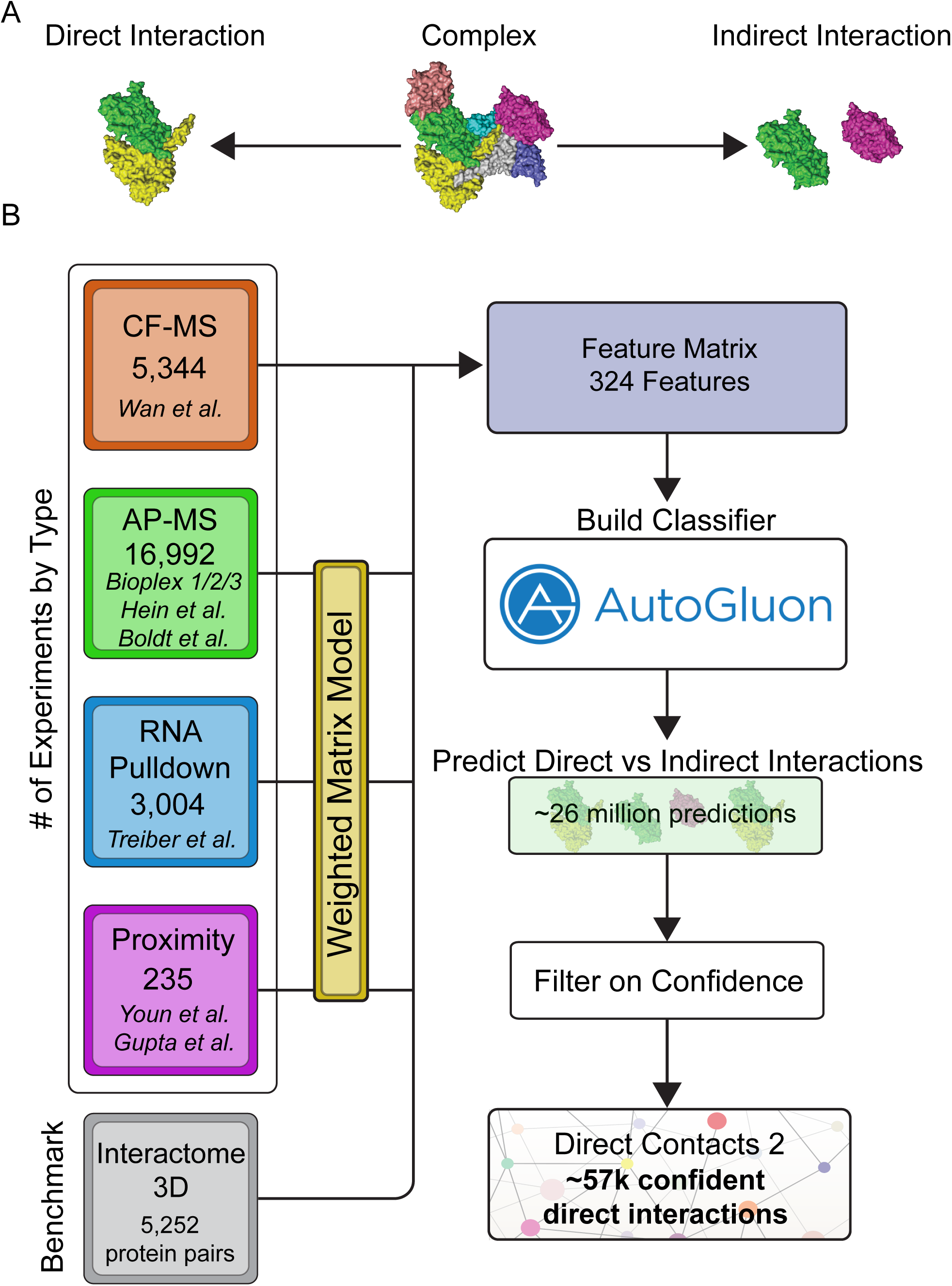
Classification of Direct and Indirect Protein Interactions Across the Human Proteome. **A**. Image displays an illustrative, multi-subunit protein complex where some directly interact (yellow and green subunits) while others indirectly interact through other subunits (magenta and green subunits). **B**. Workflow of our machine learning pipeline to integrate high-throughput proteomics data to discriminate between direct and indirect protein interactions. Data from multiple sources was used to train an ensemble machine learning model and then applied to ~26 million human protein pairs. The final network represents high confident direct interactions.

Recent advances in computational protein structure prediction now allow for near atomic structural modeling of proteins and protein complexes using sequence alone^6–9^. AlphaFold2 and RosettaFold have been applied to predict structures of protein protein interactions across *E.coli*, yeast, and human proteomes^10–14^. However, current limitations in computational resources prevent scaling these methods to all pairs in the proteome (e.g. 200 million theoretical protein pairs in the human interactome). Non-negligible false positive rates of structural modeling algorithms also limit scalability. To limit the search space of true interactors, Burke et al. applied AlphaFold2 to high-confidence predictions from hu.MAP2.0^15^, a network of co-complex interactions trained on high throughput mass spectrometry data, and showed an enrichment of high scoring structure predictions over random pairs^14^. Interestingly, while hu.MAP2.0 showed enrichment for high-confidence structural models, the hu.MAP2.0 model was originally designed to classify pairs that co-complex, regardless of whether or not they interact directly. We reasoned that a machine learning model designed specifically to differentiate direct from indirect protein interactions will generate an accurate network of directly interacting pairs and ultimately enrich for high-confidence computationally derived protein interaction structural models.

Here, we aim to identify direct physical interactions among proteins in human cells. Using data from over 25,000 mass spectrometry experiments, we developed DirectContacts2, a machine learning classifier that discriminates between direct and indirect protein interactions in protein complexes. In previous work, we showed thousands of co-fractionation mass spectrometry (CF-MS) experiments could aid in the identification of physical protein contacts^16,17^. Here we drastically improve on prior efforts in terms of accuracy and scale, by incorporating CF-MS data with affinity purification mass spectrometry (AP-MS)^18–22^, RNA pulldown experiments^23^, and a reanalysis of proximity labeling experiments^24,25^ to extract proteomic indicators of physical interactions. We built a machine learning classifier to integrate these features into a single probabilistic score which we then applied to ~26 million human protein pairs. We show our results outperform our previous DirectContacts model, hu.MAP2.0, hu.MAP3.0, and the HuRI reference map of human binary protein interactions^26^ on a benchmark of Protein Data Bank (PDB)-curated direct protein interactions. We then show our results boost the ability of identifying highly accurate AlphaFold2/AlphaFold-Multimer predictions when applying these structure prediction tools across the human proteome. We extend this result to ~2,500 new AlphaFold3 models. Finally, we demonstrate the ability to use DirectContacts2 predictions to guide structural model predictions for disease-associated complexes and place human disease mutations in structural context using a case study of a complex implicated in ciliopathy.

## Results

To construct our network of direct protein protein interactions we integrated 25,575 publicly available mass spectrometry experiments from high-throughput proteomic efforts using a machine learning framework trained on experimentally derived direct interactions found in the PDB. Our approach integrates orthogonal experimental datatypes, a strategy that has been successful in the past for building co-complex protein interaction networks such as hu.MAP^15,27,28^. Here, we integrate co-fractionation mass spectrometry (CF-MS) data from Wan et al^17^, affinity purification mass spectrometry (AP-MS) data from Bioplex 1,2,3^18–20^, Boldt et al.^22^, Hein et al.^21^, and other high-throughput proteomics datasets^23–25^. CF-MS experiments involve the biochemical separation of native protein complexes followed by mass spectrometry identification of proteins in collected fractions. We have shown previously^16,17^ that proteins in direct physical contact are more likely to coelute in a CF-MS experiment. These direct interactions can therefore be discriminated from indirectly interacting proteins by the higher similarity coefficient (e.g. Pearson correlation) of their elution profiles. In total, we construct features from 5,344 CF-MS fractions (Figure 1B), comparing nearly 1 million protein pairs (Supplemental Figure 1A), and covering 10,396 proteins (Supplemental Figure 1B).

AP-MS experiments involve isolating native protein complexes followed by a series of washes and identification by mass spectrometry. We therefore expect proteins that physically interact with the bait protein to show higher abundance in AP-MS experiments than indirectly interacting proteins. In addition to evaluating AP-MS experiments individually, we previously showed that analyzing AP-MS data collectively, using a weighted matrix model (WMM), improves co-complex interaction prediction^15,27,28^. We implement the WMM by applying the hypergeometric test to a pair of proteins across the many thousands of AP-MS experiments. In essence, the WMM asks how often a pair of proteins are seen together in the same set of experiments and whether they are seen more often than random chance. In this application, we expect pairs seen in the same set of experiments far more than random chance to be more likely to directly interact, thus allowing us to discriminate between direct and indirect interactions. This is likely due to directly interacting proteins being pulled down together in more experiments than proteins that are indirectly interacting. In total, our model incorporates data from 16,992 AP-MS experiments (Figure 1B), covering >22 million protein pairs (Supplemental Figure 1A) and 17,100 proteins (Supplemental Figure 1B).

In addition to AP-MS, we also incorporate WMM features derived from proximity labeling and RNA pulldown experiments into our model. Each individual experiment is expected to have limited signal for direct interactions. However, when we apply the WMM method to these datasets, they do provide statistical evidence for direct interactions (see section Signal from Individual Features). Specifically, we applied the WMM to 235 proximity labeling experiments^24,25^ and 3,004 RNA Pulldown experiments^23^ (Figure 1B). In total, we integrate 25,575 mass spectrometry experiments, encompassing data for ~26 million protein pairs (Supplemental Figure 1A) and 17,233 proteins (Supplemental Figure 1B), into a machine learning framework to build a classifier that discriminates between direct and indirect protein interactions.

### Integration of Proteomics Data Improves Discrimination Between Direct and Indirect Interactions

To build a network of directly interacting proteins, we reasoned that the integration of many orthogonal features derived from mass spectrometry experiments will increase our ability to discriminate between direct and indirect protein interactions. We integrated these features using AutoGluon, a flexible machine learning model selection framework^29^. AutoGluon fits multiple models to training data, including RandomForests and neural networks, and evaluates each model’s accuracy using cross-validation. AutoGluon also utilizes ensemble methods such as stacking and boosting to further increase accuracy and manage overfitting. We input 324 features from the mass spectrometry experiments, described above, into the AutoGluon framework labeled with 2,177 known directly interacting pairs of proteins and 100,707 known indirectly and non-interacting (i.e. non-co-complexing) pairs of proteins (Supplemental Table 1 & 2). Labels were derived from the Interactome3D database^30^ which curates pairs of physically interacting proteins from three dimensional structures in the PDB^31^. We partitioned a leave-out set of protein pairs that were excluded from training to be used for evaluation (Supplemental Table 3 & 4). See below and methods section for details.

Using AutoGluon, we first trained 13 base models on our labeled proteomics data. AutoGluon then trains a WeightedEnsemble_L2 model using all 13 base models (Supplemental Table 7). Only the LightGBM base model resulted with a weight > 0 after training the final WeightedEnsemble_L2 model. We then applied this final classifier to 25,992,007 protein pairs (Supplemental Table 8). The DirectContact2 score shows a bimodal distribution (Supplemental Figure 2A), with most pairs having low DirectContacts2 scores while a substantial peak is seen with high DirectContacts2 scores. This observation is consistent with our expectation that most protein pairs do not interact directly, while a subset of pairs that interact directly are identified by our model. Based on this distribution, and that of Supplemental Figure 2B, we selected two thresholds to be considered confident (DirectContacts2 score ≥ 0.7) and highly confident (DirectContacts2 score ≥ 0.9). Using these thresholds, we identify 57,494 pairs as confident and 24,328 as highly confident (Supplemental Table 9).

### DirectContacts2 Network Outperforms Previous Networks on Leave Out Set of Physically Contacting Protein Pairs

We next set out to evaluate our DirectContacts2 predictions on a large leave-out set of complexes with solved three-dimensional structures. Specifically, as done for our training set, we used a set of directly contacting protein pairs from the Interactome3D database as a leave-out test set to evaluate the network’s performance. We only included complexes of size 5 subunits or more in our test set to eliminate the possibility of trivial cases where every subunit in a complex is directly interacting (e.g. dimers). All other pairs of proteins within a complex without a direct interface are labeled as indirect pairs (i.e. negatives). We excluded the negative set pairs from separate complexes to provide a rigorous assessment of our model. In other words, we evaluate our model on its power to discriminate direct and indirect interactions within the same complex. Additionally, we ensure the test set is disjoint from the set used for training. This resulted in a 40:60 split of direct vs indirect pairs in the test set.

Supplemental Figure 2B shows the distribution of DirectContact2 scores for positively labeled (i.e. directly interacting) and negatively labeled (i.e. indirectly interacting) pairs from our leave-out test set. We see significant separation between the two distributions where positive labeled pairs are scored significantly higher than negatively labeled pairs (Mann Whitney p-value: 4e-18), indicating our model’s ability to distinguish the two. Performing a precision-recall analysis using this same leave-out set, we observe that our DirectContacts2 predictions outperform the hu.MAP2.0 network, the hu.MAP3.0 network, the HuRI yeast2hybrid network, as well as our previous DirectContacts model (Figure 2A). The Random curve represents the background probability of identifying a positive label with ~40% of direct pairs in the test set. Figure 2B shows the area under the precision-recall curve (AUC) for each network, and again demonstrates that DirectContacts2 outperforms other networks. In a more global precision-recall analysis that included additional negative pairs from separate complexes, DirectContacts2 still achieved the best performance of all models tested (Supplemental Figure 2C).

**Figure 2:**
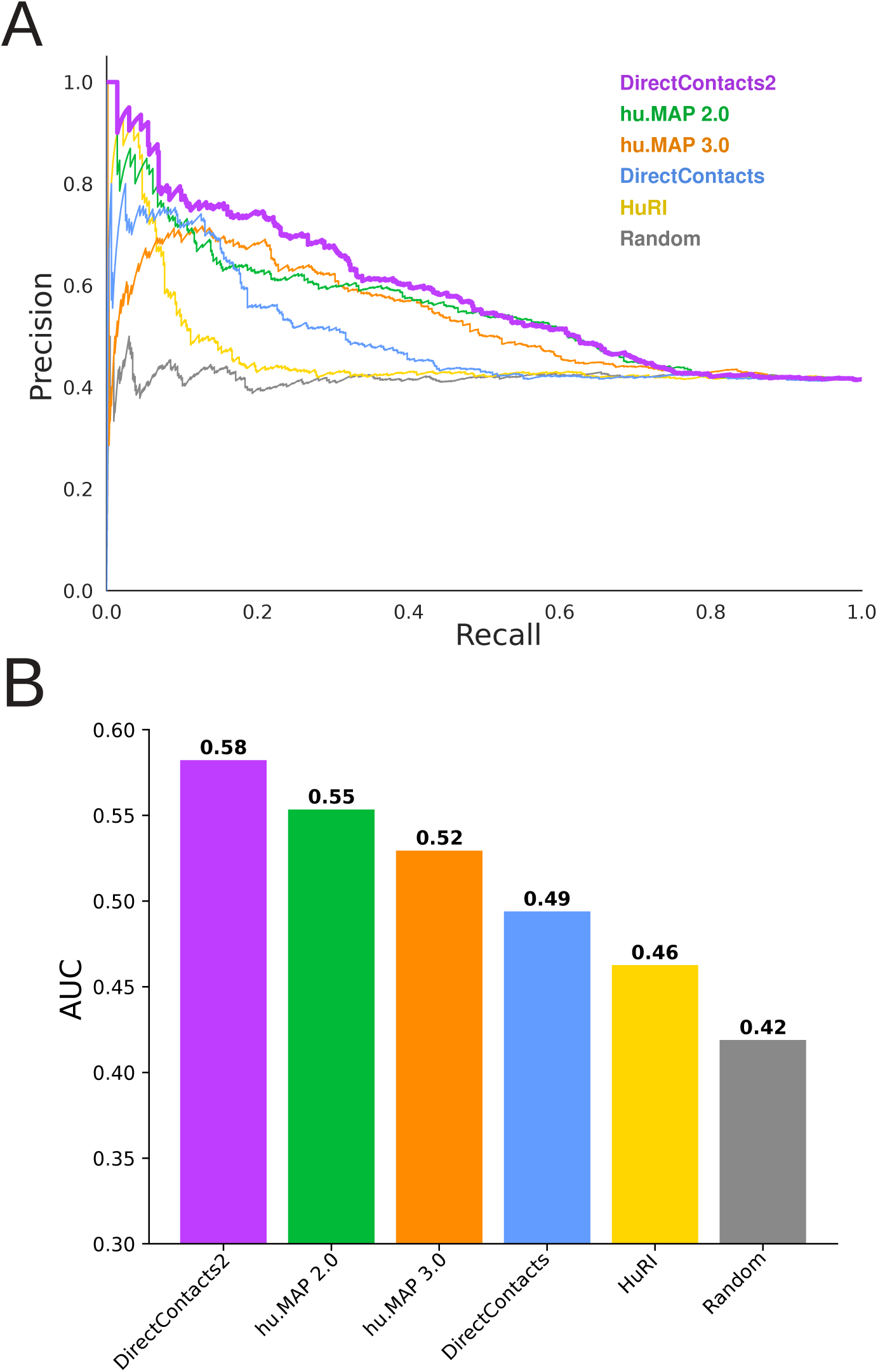
DirectContacts2 is Highly Accurate Compared to Alternative Networks. **A.** Precision recall analysis shows the DirectContacts2 model (purple) outperforms the HuRI network (yellow), hu.MAP2.0 (green), hu.MAP3.0 (orange), and the original Direct Contacts model (blue). A random shuffling of pairs is shown in gray. Analysis is performed on the benchmark leave-out set of direct (positive) and indirect (negative) protein interactions. **B.** Area under the precision recall curve bar plot shows quantitatively that DirectContacts2 outperforms other models and networks.

### Investigation of DirectContact2 Predictions of a Multisubunit Complex: chaperonin CCT-Prefoldin complex

We next look at our model’s ability to discriminate between direct and indirect interactions on a specific example of a multisubunit complex from our leave out set of test complexes. In Figure 3A, we show the three-dimensional structure of the chaperonin CCT-Prefoldin complex (PDBID: 6NRB^32^) with each protein chain colored differently. DirectContacts2 makes predictions for 46 pairwise interactions within the complex, 23 direct and 23 indirect. Figure 3B shows the DirectContacts2 confidence score distributions for these two sets of predictions, in which there can be seen a clear distinction between the known positives and negatives, with direct interactions receiving higher scores by the model. The two distributions are significantly distinct using the Mann-Whitney test (p-value = 0.002). Notably, of the top 8 highest scoring pairs of proteins from the chaperonin CCT-Prefoldin complex, seven are true positives and only one is a false positive (Figure 3C). This aligns with our global analysis for the complete test set (Supplemental Figure 2B), and further confirms that our DirectContacts2 network successfully discriminates between direct and indirect protein interactions within specific examples on which it was not trained.

**Figure 3:**
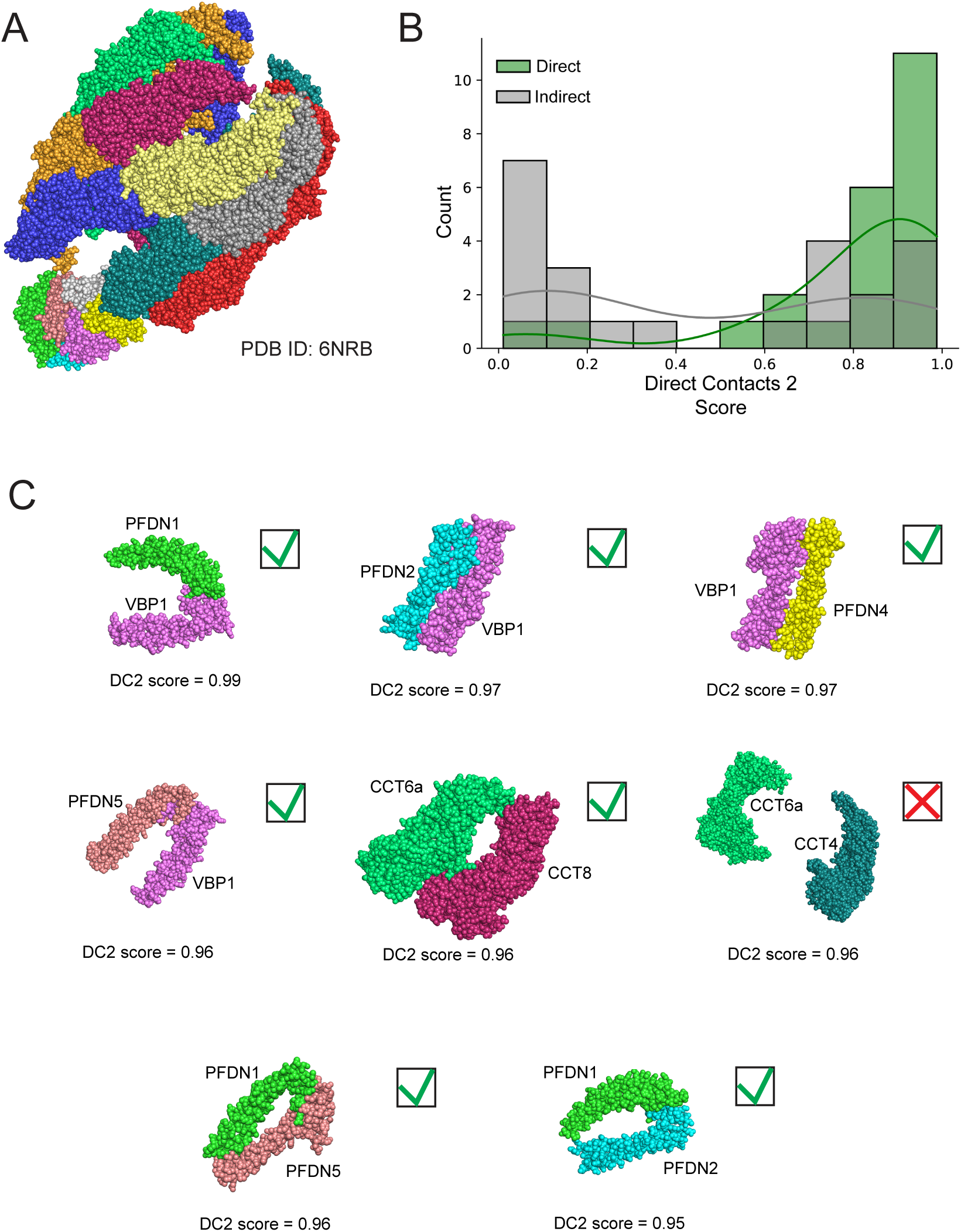
DirectContacts2 Accurately Identifies Physical Interactions of the chaperonin CCT-Prefoldin complex. **A.** Three-dimensional structure of the chaperonin CCT-Prefoldin complex from test leave out set (PDBID: 6NRB). **B.** Distributions of DirectContact2 scores for directly interacting pairs (green) and indirectly interacting pairs (gray) from chaperonin CCT-Prefoldin complex. The directly interacting pairs have a right shifted distribution with higher DirectContact2 scores Mann-Whitney u-Test statistic=406.0, p-value=0.002. **C.** Eight highest scoring protein pairs within the chaperonin CCT-Prefoldin complex. Seven of the 8 high scoring pairs correctly predict direct interactions. Green checkmark = true positive, red X = false positive.

### Enrichment of High Confidence Multichain AlphaFold2 Models in DirectContact2 Predictions

Since the advent of AlphaFold2/AlphaFold-Multimer and RoseTTAFold^6,8,9^, several groups have applied these new capabilities to modeling protein interactions within the human proteome^10,13,14^. Due to the scale of all possible pairwise interactions across a proteome, (N^2 where N is the number of proteins in the proteome), applying computational structural modeling to all proteome pairs is currently infeasible, and protein pairs must therefore be prioritized.

A recent study by the Elofsson and colleagues applied their FoldDock algorithm^14^ based on AlphaFold2 to generate structural models for 10,207 high-ranking hu.MAP 2.0 co-complex interactions in addition to 55,773 interactions from HuRI, derived from high-throughput yeast2hybrid experiments^14,15,26^. Using a score metric (pDockQ) that evaluates the quality of the modeled interaction, they observed an enrichment of pairs with highly confident pDockQ scores (≥0.5) in our previous hu.MAP2.0 network. We therefore theorized that our DirectContacts2 network should achieve greater enrichment for models with high pDockQ values due to its more stringent focus on direct physical interactions rather than just co-complexing interactions compared to our hu.MAP networks, which are a mix of direct and indirect interactions.

To test if the DirectContacts2 model’s predictions are enriched for quality of structural models over previous networks, we utilize an expanded set of 277,807 AlphaFold models^13,14^. To evaluate the networks on a fair basis, we calculate the fraction of each network’s most confident set of predictions with high-confidence AlphaFold models (pDockQ > 0.5) for increasing sizes of top-ranked pairs. In Figure 4A, we observe 32% of the top 2,500 (DirectContacts2 score > ~0.99) predictions contain high-confidence AlphaFold models. This outperforms hu.MAP3.0 (29%), DirectContacts1 (25%), hu.MAP2.0 (17%), and HuRI (15%). This trend remains consistent for all greater numbers of top predictions (Figure 4A). Using a benchmark of large heteromeric protein complexes, the Elofsson study highlighted that 38% of known directly interacting pairs resulted in having highly confident pDockQ scores (>0.5), suggesting an upper bound^14^. Comparing this with our result in Figure 4A suggests that our DirectContact2 model is near the upper bound of performance. Additionally, we perform the same rank-based analysis of pDockQ score enrichments between networks at the minimum threshold for confidence (pDockQ ≥ 0.23)^33^, at which the relative enrichment of DirectContacts2 pairs for interface quality over other networks becomes even more substantial (Supplemental Figure 2D).

**Figure 4:**
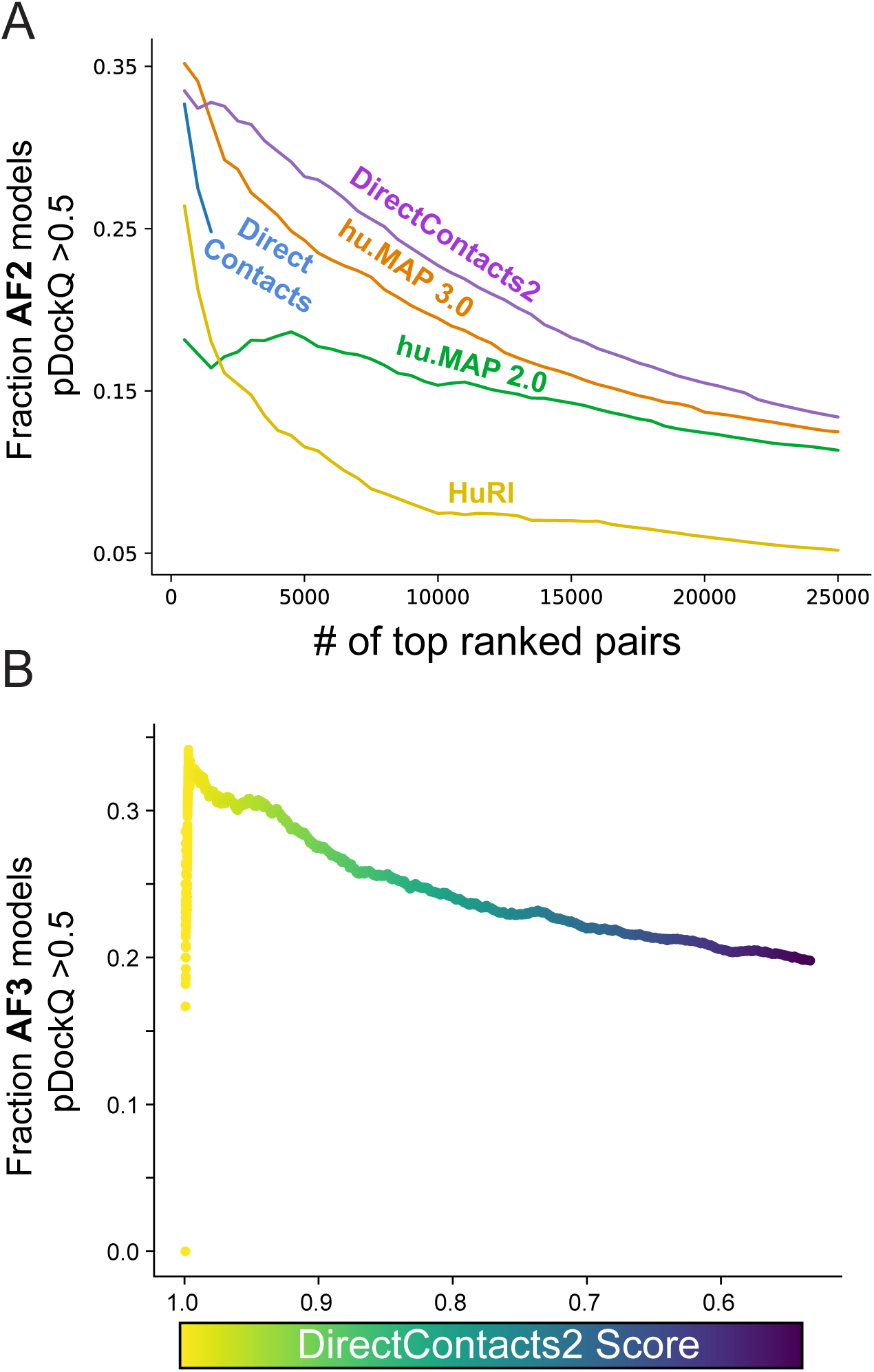
DirectContacts2 Enriches for Confident AlphaFold Protein Interaction Structural Models. **A.** Comparison of enrichment for high-confidence pair interfaces out of the set of top confident pairs for DirectContacts2 and other networks. For each set, the fraction of high-confidence interfaces (pDockQ ≥ 0.5) out of their top predictions was calculated for increasing numbers of top predictions. **B**. The fraction of AlphaFold3 protein interaction models with high-confidence pDockQ pairs.

### Generation of AlphaFold3 models from high confidence DirectContacts2 predictions

While we observed a general enrichment for higher quality interfaces in existing AlphaFold structural datasets, we wanted to evaluate the practical utility of our DirectContacts2 score towards guiding new structural predictions. We produced AlphaFold3 models for 2,573 pairs that our model predicted to have a DirectContacts2 score ≥ 0.5, and then assessed their pDockQ scores. We see in Figure 4B a similar trend to our analysis above, in which high-confidence DirectContacts2 pairs exhibit a greater enrichment for structural quality when modeled with AlphaFold3. A table of AlphaFold3 predictions and their scoring metrics can be found in Supplemental Table 10.

### Intermolecular Cross-links are enriched in DirectContact2 Network

Mass spectrometry cross-linking (XL-MS) experiments have the ability to identify directly interacting pairs of proteins from *in vivo* experiments. Since no XL-MS experiments were used in the training of DirectContacts2, cross-linking provides an additional independent means of benchmarking the DirectContacts2 network. Using the set of 5,401 intermolecular cross-linked protein pairs identified in Wheat et al.^34^, we observe a substantial overlap between the DirectContacts2 network’s most confident pairs and the crosslink-derived network when compared to random pairs of proteins (Z Score = 65.2) (Figure 5A). Since the DirectContacts2 network was constructed using the same feature matrix as hu.MAP3.0, we asked whether this enriched overlap is due to the direct physical nature of DirectContacts2 as opposed to simply co-complex interactions. To answer this, we perform the same overlap analysis between the hu.MAP3.0 network’s top pairs and the XL-MS dataset. Comparing the two networks on the basis of their resulting Z Scores, we see an increase in overlap of our DirectContact2 predictions with protein crosslinks when compared to hu.MAP3.0 high ranking pairs (Z Score 65.2 vs 48.7) (Figure 5A), demonstrating an enrichment for direct physical interactions above co-complex interactions.

**Figure 5:**
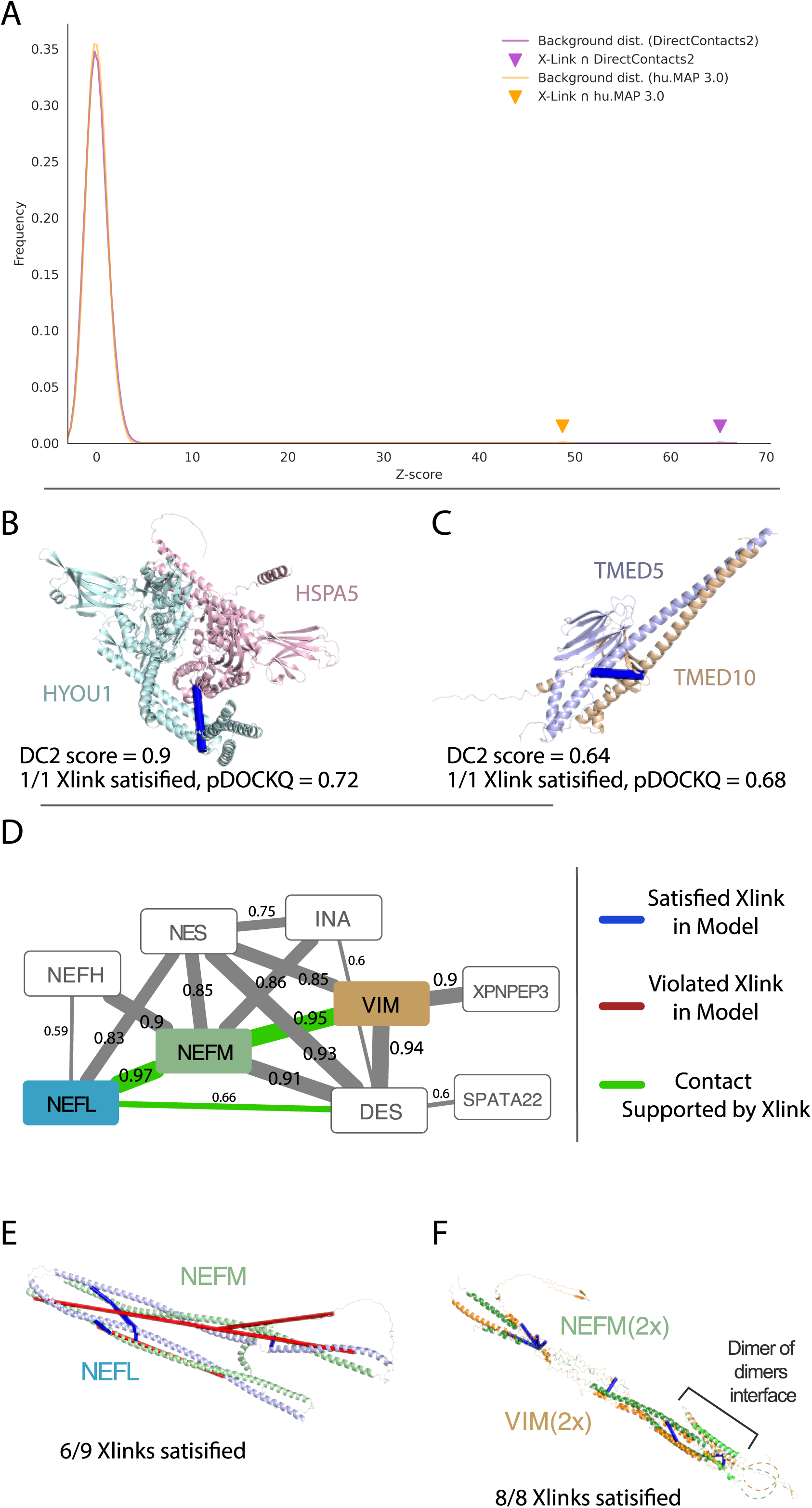
Crosslinking supports DirectContact2 predictions. **A.** Overlap of crosslinking pairs from Wheat et al. with DirectContacts2. DirectContacts2 Z-score (purple triangle) is calculated by comparing the overlap of the top 57,494 DirectContacts2 pairs (DirectContacts2 score > 0.7) with crosslinked pairs to a distribution of overlap of random pairs and crosslinked pairs (purple histogram). DirectContacts2 Z-score is substantially increased in comparison to the top 57,494 hu.MAP3.0 pairs’ overlap with crosslinked pairs (orange triangle). **B-C**. Examples of high-confident DirectContacts2 predictions (HSPA5-HYOU1 and TMED5-TMED10) supported by both AlphaFold2 models and crosslinks. Both examples show crosslink is satisfied (blue) by the structural model. **D.** DirectContacts2 network of intermediate filament complex (HuMAP2_01819). Multiple direct edges are supported by experimental crosslinks (green). Value on each edge represents DirectContacts2 score. **E.** AlphaFold3 model of highest confident DirectContacts2 prediction (NEFL-NEFM) from intermediate filament complex (panel **D**). The majority of experimental crosslinks are satisfied (6 satisfied = blue, 3 violated=red). **F**. AlphaFold3 model of second highest confident DirectContacts2 prediction (NEFM-VIM) from intermediate filament complex (panel **D**). Dimer of dimers model shows all 8 experimental crosslinks are satisfied.

We next focus on individual examples of interactions and show agreement between DirectContacts2 predictions, XL-MS cross-links, and AlphaFold2 models. Figure 5B shows the pair HSPA5 and HYOU1 which has a highly confident DirectContact2 score of 0.9 and a single experimental MS cross-link HSPA5 Lys370 and HYOU1 Lys883. Although there is no experimental structural coverage of these two proteins, an AlphaFold2 model of HSPA5 and HYOU1 from Burke et al.^14^ shows the MS cross-link between these residues is satisfied (pDockQ = 0.72). Similarly, the pair of proteins TMED5 and TMED10 was predicted to directly interact (DirectContacts2 score = 0.64) and has an experimental MS cross-link (TMED5 Lys169 and TMED10 Lys170) that is satisfied by an AlphaFold2 model (pDockQ = 0.68, Figure 5C).

While this prediction is below our confidence threshold of 0.7, the example highlights that a combination of orthogonal data, such as XL-MS, can provide supporting evidence for direct interactions. These examples demonstrate alignment between DirectContacts2 and cross-linking data and further points to the ability to build high-quality structural models for protein complexes using DirectContacts2 as a guide.

We next turn to a larger complex with high confident DirectContacts2 predictions which does not have an experimental structure. The intermediate filament complex (HuMAP2_01819) consists of several filament subtypes but knowledge of their true direct interactions is limited (Figure 5D). Intermediate filaments are involved in cytoskeleton construction and their disruption leads to several human diseases^35^. Our DirectContacts2 model identifies confident direct interactions among several pairs of subunits in the complex. In particular, we observe the NEFL-NEFM and NEFM-VIM pairs to have the highest confidence scores within the complex, 0.97 and 0.95 respectively. We identify XL-MS crosslinks^34^ supporting these interactions, providing additional evidence for these direct interactions (Figure 5D). We next modeled the two subcomplexes, NEFL-NEFM and NEFM-VIM, using AlphaFold-multimer to gain structural intuition of the interactions. Both resulting models produced two elongated coiled coil structures linked by a short loop region (Figure 5E and 5F). Supporting these structures, recent work by Jänes et al. have generated AlphaFold models for both pairs, NEFL-NEFM and NEFM-VIM, both of which produced confident pDockQ values of 0.72^13^. We next map XL-MS cross-links from the Wheat et al set onto our NEFL-NEFM AlphaFold-multimer model and observe that the model satisfied 6 out of 9 experimental cross-links. Intriguingly, the residues of one crosslink in the NEFL-NEFM model, NEFL:Lys91 - NEFM:Lys271, are located at opposite ends of the structure with a distance of ~212Å. This suggests the crosslink may be satisfied by a higher order oligomerization of the NEFL-NEFM subcomplex.

We next evaluated the NEFM-VIM subcomplex on a similar basis and constructed an initial NEFM-VIM model that satisfied 6 out of 8 crosslinks, with the 2 violated crosslinks on opposite ends of the structure (NEFM:Lys259 - VIM:Lys120 : ~211Å, NEFM:Lys,271 - VIM:Lys120 : ~198Å). McCafferty et al. recently showed large distances between cross-linked residues agree with oligomerization models of proteins^36^. Furthermore, intermediate filaments are known to form higher order structures^37^, so we theorized that these unsatisfied cross-links may point to sites of oligomerization. Using a structural model built from the expected overlapping segments, we constructed a higher order model of NEFM and VIM. When modeled as a higher order oligomer, the remaining unsatisfied cross-links between NEFM and VIM were satisfied (Figure 5F). Taken together, we show using DirectContacts2 predictions with complimentary structural data such as XL-MS aids in the modeling of protein complex structures.

### DirectContacts2 guides the structural model of a human ciliopathy complex

OFD1 - PIBF1 - FOPNL(CEP20,FOR20) - KIAA0753 form a complex whose members localize to the distal ends of basal bodies and are important for proper ciliogenesis^38,39^. Our hu.MAP2.0 network indicates that these four genes are components of a larger complex (HuMAP2_00328), the Orofacial Digital Syndrome (OFDS) complex. Of the pairwise interactions in this complex, OFD1-FOPNL and FOPNL-KIAA0753 exhibit the highest DirectContact2 scores of 0.91 and 0.98 respectively. Mutations in these genes are causative of severe clinical phenotypes through the disruption of cilia formation^40–43^. However, there is currently no experimental structural data to interpret these mutations in the broader structural context of the proteins’ molecular interactions. We therefore used our DirectContacts2 predictions to direct AlphaFold-multimer modeling of the three-dimensional structure of OFD1-FOPNL and FOPNL-KIAA0753 (Figure 6A and 6B). The interfaces of the models resulted in highly confident pDockQ scores of 0.63 and 0.57 respectively, indicating accurate modeling of the dimers. The Predicted Aligned Error (PAE) plots produced by AlphaFold-multimer (Figure 6C and 6D) further suggest high global confidence of the structure. The FOPNL-KIAA0753 model was consistent with previous work that identified the C-terminus of KIAA0753 as being necessary and sufficient for the binding of FOPNL^38^.

**Figure 6:**
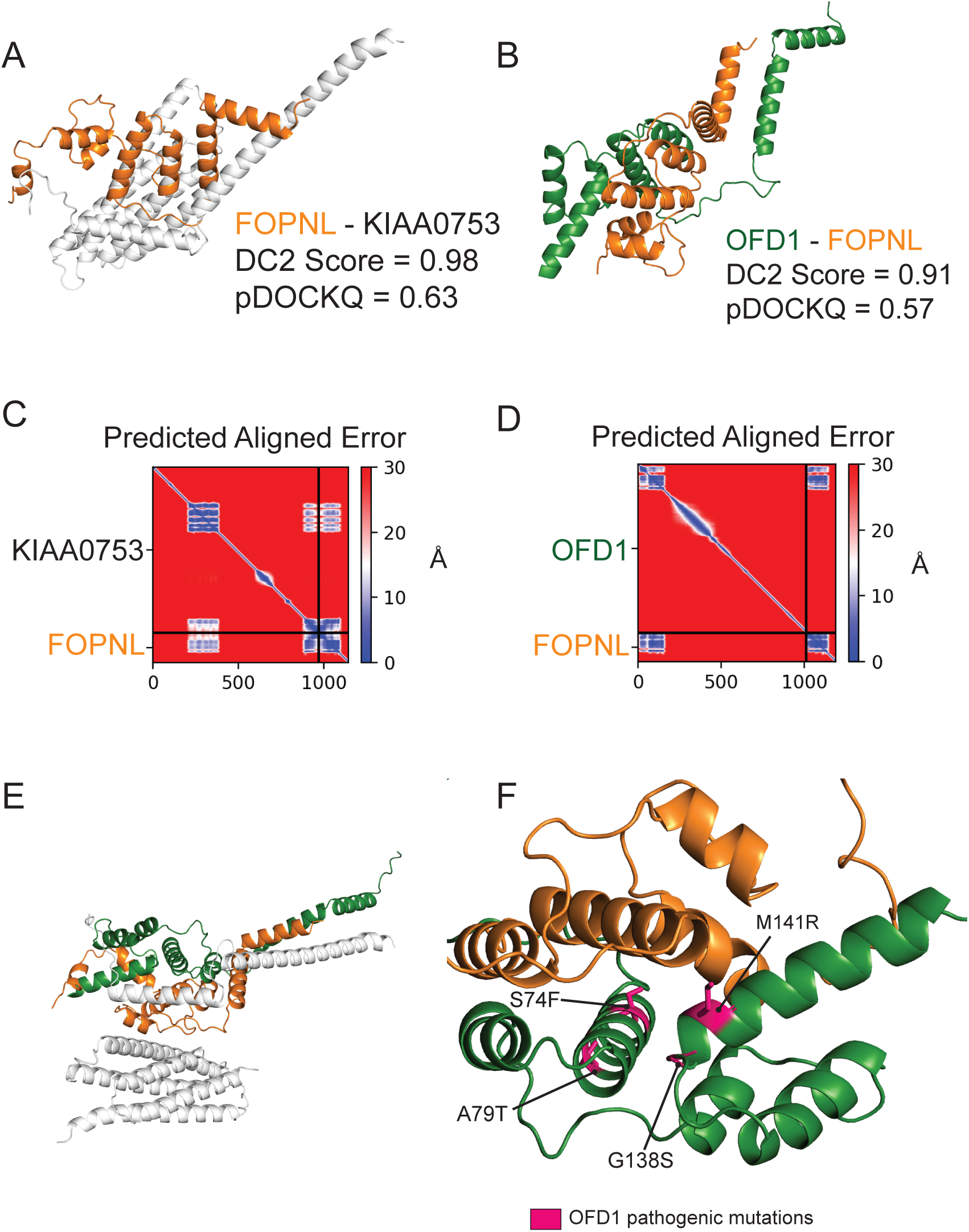
Direct contact network of a ciliary protein complex informs structural modeling. **A.** AlphaFold-Multimer structure prediction of FOPNL-KIAA0753, highlighting area around the interface. **B.** AlphaFold-Multimer structure prediction of OFD1-FOPNL, highlighting area around the interface. **C.** Predicted Aligned Error graph of the FOPNLKIAA0753 AlphaFold-Multimer prediction. **D.** Predicted Aligned Error graph of the OFD1-FOPNL AlphaFold-Multimer prediction **E.** Predicted interface of the OFD1-FOPNLKIAA0753 complex by overlapping FOPNL chains. **F.** OFD1 disease mutations (pink) cluster at the OFD1-FOPNL interface.

Additionally, we observed another region of KIAA0753 (aa 206-380) that formed a substantial interface with FOPNL, with an average per-residue pLDDT value > 80. To construct a higher order structure of the complex, we aligned the two structure models on their common subunit (FOPNL) and observed distinct interfaces between the three subunits (Figure 6E). The resulting model placed four sites of OFD1 mutation (i.e. rs312262812, rs312262814, rs312262827, rs886039860) at the interface of FOPNL (Figure 6F), strongly suggesting that the disruption of this interface is responsible for the downstream functional defect giving rise to a clinical phenotype.

### Signal From Individual Features

Machine learning algorithms excel at integrating multiple lines of evidence from orthogonal sources towards building powerful classifiers. We next asked which features provide the most signal in discriminating between direct and indirect protein interactions. In Supplemental Figure 3, we show the top 20 features ranked based on the calculated area under the precision-recall curve (AUPRC) evaluated on the benchmark used to train our classifier. We have shown in previous work that CF-MS features could be used as powerful predictors of physical interactors^17^. Here, we see again that CF-MS data provides strong predictive signals of interaction, with 16 of the top 20 features in the DirectContacts2 model being CF-MS features. We also see AP-MS features exhibiting strong predictive capability, including the Bioplex3 feature ‘uPeps’, which represent the quantification of the number of unique peptides for a given prey protein. This feature originates from two cell types in the Bioplex3 dataset, HCT116 cells and HEK293T, and perform very well as discriminators of direct and indirect interactors.

Additionally, we observe WMM features among the top features suggesting that our WMM can discriminate between direct and indirect protein interactions as well.

Somewhat unexpectedly, WMM features calculated from proximity labeling experiments^24,25^ had very high performance under the AUPRC measure (Supplemental Figure 3). Due to the diffusible nature of labeling in proximity labeling techniques, features from individual proximity labeling experiments are not expected to be valuable discriminators between direct and indirect interactions. In fact, we see a limited number of pairs in our training set (9 total pairs) that have evidence from proximity labeling experiments (non-WMM features) where one subunit is a bait and the other is the prey. Furthermore, several bait-prey proximity labeling features representing data from an individual experiment are automatically removed from consideration by the AutoGluon framework during training due to their limited signal. While an individual proximity labeling experiment doesn’t appear to have signal for this application, the WMM which statistically analyzes many experiments collectively shows great statistical power for discriminating between direct and indirect protein interactions (neg_ln_pval_youn_hygeo_gt4 and neg_ln_pval_cilium_hygeo features, Supplemental Figure 3).

To investigate this further, we return to the example of the CCT-Prefoldin complex interactions. The Prefoldin complex is a chaperone that physically engages with the TRiC/CCT complex to prevent misfolding and helps maintain proteostasis. As described above, a recent cryo-EM structure of Prefoldin in complex with the TRiC/CCT provides us with a specific test case of our WMM feature^32^. Supplemental Figure 4A shows a heatmap of 20 proximity labeling experiments where at least one Prefoldin subunit is identified above an abundance threshold (see Methods, ≥4 average PSMs). We observe two distinct regions that suggest individual Prefoldin subunits are isolated together without other subunits. Specifically, we see PFDN5 and PFDN3 (highlighted box in yellow) as well as PFDN3 and PFDN2 (highlighted box in purple) are seen in several shared experiments, pointing to their likelihood to directly contact each other. Notably, we do not see PFDN1 and PFDN6 together without the other subunits of the complex, which is consistent with their spatial separation and lack of a direct interface as shown in the cryo-EM structure of the Prefoldin complex (PDBID: 6NRB) (Supplemental Figure 4B).

We next evaluate our calculation of the hypergeometric test on these data as we expect it to more finely discriminate direct from indirect interactions. The hypergeometric test is a rigorous statistical test which asks how often a pair of proteins are seen together in the same set of experiments and are they seen more often than random chance. As described above, this is the central concept of the WMM calculation. To discriminate between direct and indirect interactions we expect the values calculated by the hypergeometric test to exhibit a bimodal distribution where indirect interactions correspond to lower scores than direct interactions. Supplemental Figure 4C shows the distribution of the −lg(Pval) calculated from the hypergeometric test for all observed pairs of Prefoldin subunits, and exhibits the expected bimodal topology (direct = higher scores, right density and indirect = lower scores, left density) with the PFDN5/PFDN3 and PFDN3/PFDN2 pairs having higher overall scores.

Comparing this distribution to the cryo-EM structure of Prefoldin, we see that PFDN3 and PFDN5 form substantial direct interfaces (Supplemental Figure 4B). Similarly, we see that PFDN3 and PFDN2 also make direct physical contact with each other in the structure, further validating our observations. Lastly, PFDN1 and PFDN6 do not interact, as shown by their locations on opposite sides of the complex, which is also quantitatively reflected in their placement within the left density of the bimodal −lg(Pval) distribution (Supplemental Figure 4C). These results show on an individual example how the WMM computed on proximity labeling experiments can discriminate between direct and indirect protein interactions, and when taken together with our results from Supplemental Figure 3, demonstrate that the combination of WMM and proximity labeling is a valuable feature for predicting direct interactions on the proteome scale.

## Discussion

Determining a comprehensive wiring diagram of the cell is an important challenge in biological research. A network of directly interacting biomolecules provides molecular mechanisms for understanding disease and molecular function. In addition, it provides a global view of how the cell is constructed, leading to functional insights of the system as a whole. Here we construct DirectContacts2, a network of directly interacting proteins, by integrating a broad array of high-throughput proteomics experiments. Our integration was based on a machine learning framework trained on directly contacting proteins identified from experimentally solved protein structures (Figure 1). We showed our network outperformed the state-of-the-art in identifying direct interactions (Figure 2) as well as enriching for high-confident AlphaFold2 models (Figure 4). Furthermore, our network shows good concordance with crosslinking data, which is an orthogonal experimental method for identifying direct protein interactions (Figure 5). Finally, we demonstrate the utility of our network by building and analyzing AlphaFold models of protein interactions without known experimental structures including ~2,500 AlphaFold3 models.

Specifically, we analyze interactions with high DirectContact2 scores, including the HYOU1-HSPA5, TMED5-TMED10, NEFL-NEFM, NEFM-VIM interactions, and demonstrate support for their direct interactions using high-confidence AlphaFold models and crosslinking data (Figure 5). Finally, we identify developmental disease genes having high DirectContact2 scores with their interactors. When modeled, these interactions result in high-confidence AlphaFold2 models that provide mechanistic structural insight into several patient mutations’ roles in known clinical phenotypes (Figure 6).

While our DirectContacts2 network is more accurate than other state-of-the-art networks, our predictions still have an appreciable amount of error, both false positives and false negatives. To discriminate between direct and indirect interactions, our method attempts to extract signals from experimental proteomics data where subcomplexes are biochemically separated from their larger assemblies. As discussed previously^17^, subcomplexes of highly stable complexes (e.g. 20S proteasome core) are rarely seen apart from the entire complex in high-throughput data making it difficult to discern direct interactions. Alternatively, low-stability and transient complexes (e.g. kinase - kinase substrate interactions) are difficult to capture with the experiments utilized here. Discriminating direct interactions within highly stable complexes and capturing transient interactions therefore continues to be a challenge in the field. Identifying separation techniques that disrupt high stability complexes or stabilize low stability complexes may help capture these elusive interactions. The DirectContacts2 network is also biased towards high-abundance complexes. The power of integrating a large compendium of experiments comes from having multiple lines of evidence to inform prediction. Low-abundance proteins may rarely be seen in high-throughput proteomics experiments and therefore this makes predictions difficult for these biomolecules. However, DirectContacts2 increases the total coverage of the human proteome over our previous DirectContacts network by ~10 times (DC2 proteins = 14,834 vs DC1 proteins = 1,504).

The DirectContacts2 network (Supplemental Table 9) will be a valuable resource for the structural, computational, and proteomics research communities. We anticipate the network will have many applications including prioritizing high-throughput computational structural modeling and identifying therapeutic targets that were once considered undruggable.

## Methods

### Compilation of mass spectrometry datasets

To generate our DirectContacts2 network we utilized a feature matrix derived from published mass spectrometry experiments where each feature quantifies the statistical evidence for a pair of proteins to interact. The feature matrix consisted of the same experiments used in Fischer et al. for the construction of hu.MAP3.0 co-complex interaction network^28^. Features representing CF-MS data from Wan et al.^16^ consist of 1-D vector comparison measures including Poisson noise, Pearson correlation coefficient, weighted cross-correlation, co-apex score, and MS1 ion intensity distance metric. CF-MS features were filtered to ensure the pair of proteins contained at least one 1-D vector comparison score > 0.5 in at least one human CF-MS experiment.

Features from Bioplex1^20^ and Bioplex2^19^ included NWD Score, *Z* Score, Plate *Z* Score, Entropy, Unique Peptide Bins, Ratio, Total PSMs, Ratio Total PSMs, and Unique:Total Peptide Ratio. Bioplex3^18^ features included the same features as Bioplex1 and 2 for both HEK293T and HCT116 cell lines. AP-MS features from Guruharsha *et al.*^44^ and Malovannaya *et al.*^45^ were represented by HGSCore value and MEMOs value (mapped to “approved”=10, “provisional”=3, and “temporary”=1) respectively. Hein et al.^21^ AP-MS data included prey.bait.correlation, valid.values, log10.prey.bait.ratio, and log10.prey.bait.expression.ratio. Features from Boldt et al.^22^ included socioaffinity index (SA*ij*), spoke model indices (S*ij*, S*ji*) and matrix model socioaffinity index (M*ij*). Proximity labeling features representing single experiments from Youn et al.^24^ and Gupta et al.^25^ include Average Spectra, Average Saint probability, Max Saint probability, Fold Change, and Bayesian FDR estimate. Proximity labeling features were ultimately removed during feature selection of the classifier training workflow as they provided no predictive signal (see below). Full feature matrix can be found at https://humap3.proteincomplexes.org/static/downloads/humap3/humap3_20220625.featmap.gz.

**Table 1:**
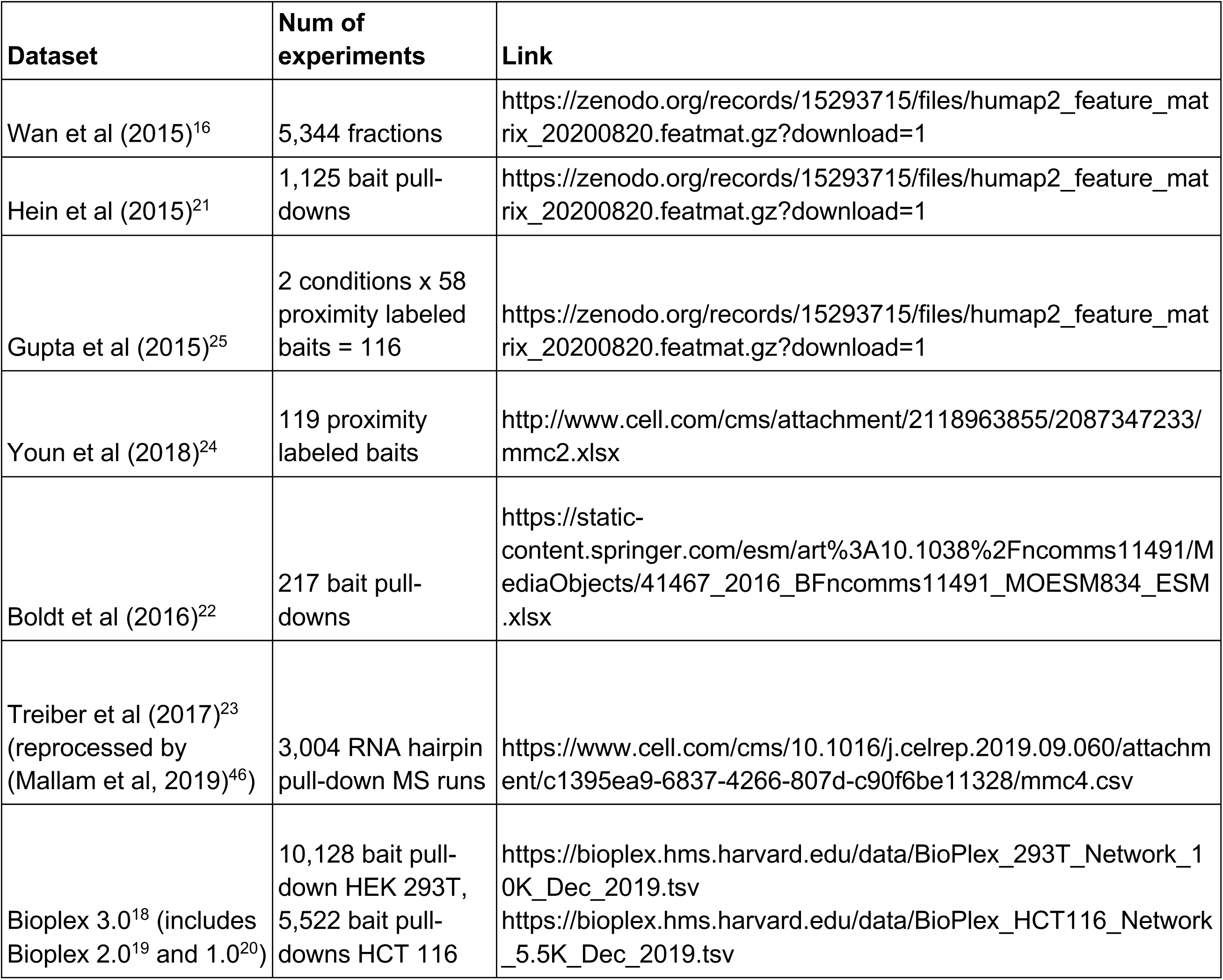
Datasets.

### Weighted matrix model calculation

Weighted matrix model (WMM) features were calculated on Bioplex1^20^, Bioplex2^19^, Bioplex3^18^ (HCT116, Hek293T, and HCT116+Hek293T combined), Treiber et al.^23^, Hein et al.^21^, Gupta et al.^25^, and Youn et al^24^. The full method was previously described in our hu.MAP^27^ study, as well as in Hart et al.^47^ The WMM is based on the hypergeometric test and asks, given two proteins A and B, whether they are seen in the same set of experiments more often than random chance using equation 1, where *k* = # of experiments both proteins, A and B, are seen together, *n* = # of experiments protein A is seen, *m* = # of experiments protein B is seen, and *N* = total # of experiments in the dataset. For each pair of proteins, a p-value is produced, which is then mapped to a negative logarithmic function (-log(p-value)), such that low p-values correspond to high-confidence interactions. In addition, we include a feature that summarizes the number of experiments each protein pair shares in a dataset (pair_count).

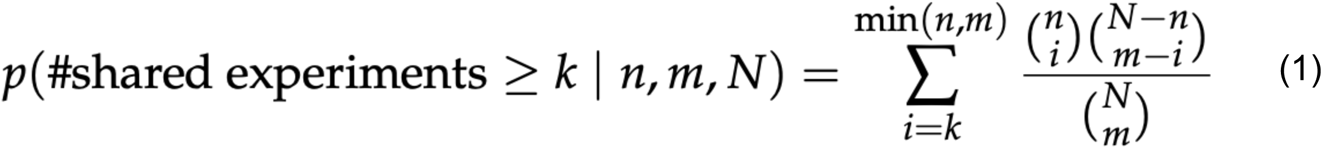

The WMM for Bioplex 3.0 was calculated for the HEK293 experiments and HCT116 experiments individually, as well as collectively for the two cell type experiments combined. We used two thresholds (Bioplex Z-score > 2 and Z-score > 4) to discern whether an individual protein was present in an experiment and then calculated the WMM (e.g. feature name: neg_ln_pval_bp3_HCT116_Z4).

### Benchmark Test and Training Dataset

Benchmark test and training datasets were derived from Interactome3D^30^ which identifies physically interacting pairs of proteins from three-dimensional structures in the Protein DataBank^31^, and collates them into ‘interactions.dat’ data tables giving high-level summaries for each pair, where an “entry” corresponds a single row in this table. To construct our benchmark, we retrieved the Interactome3D (Ver. 2020_05) *Homo sapiens* table, and then filtered for entries with associated experimental PDBs (i.e. labeled ‘structure’ in type column). We first split the Interactome3D entries based on their parent complexes (defined by PDB id). The complexes were then clustered into non-overlapping sets using a greedy approach, where cluster overlap was defined as a Jaccard score > 0 between their complex’s constituent proteins. This resulted in 5 clusters of complexes whose constituent pairs were completely unique to that cluster. The largest cluster was used as our training set, and the smaller four were combined and used as our leave-out set. See Supplemental Table 6 for a comprehensive list of positive test and training pairs.

To label our feature matrix during training, we only considered protein pairs from complexes with ≥ 3 subunits to avoid trivial cases of dimers, which are guaranteed to only have a single, direct interaction. Negative labels in the training set were defined as all pairs of proteins in the set not labeled as positive, including co-complexing pairs that do not share an interface, as well as pairs from different complexes. To gain better insight into true performance, we constructed a more rigorous leave-out test set, which included only complexes with ≥ 5 subunits. This nearly eliminates the trivial cases where all subunits in a complex share direct interfaces with all others. Also, labels of test negatives were defined to be only co-complexing protein pairs that do not share an interface within a complex. This contrasts with the training set, as it excludes negatives between separate complexes, and, therefore, is more stringent of a test. Finally, we only considered PDB structures with ≥80% sequence coverage in the structure. This ensures that protein pairs with limited structural coverage are truly indirect interactions, rather than potentially interacting through unresolved protein domains not reflected in their structures. In total, the training set consisted of 2,177 positive labels from 595 complexes of which 1,472 had entries in our feature matrix and 100,707 negative labels of which 47,246 had entries in our feature matrix. The test set consisted of 638 positive labels from 50 complexes of which 572 had entries in our feature matrix and 898 negative labels of which 785 had entries in our feature matrix. For a complete list of entries, see Supplemental Tables 1-6.

### Machine learning framework to discriminate between direct and indirect interactions

To train our classifier, we used AutoGluon’s Tabular Predictor with a binary classification scheme^29^. For training, we used our feature matrix (described above) of 324 experimentally derived features representing proteomic evidence of direct physical interaction. Labels for training were derived from Interactome3D as described above, and we set AutoGluon’s evaluation function to be ‘accuracy’, using default values for all other parameters. AutoGluon’s fit function performs cross-validation training and hyperparameter optimization for 13 different classifier architectures (Supplemental Table 7). Within Autogluon’s automated workflow, several features were deemed insufficiently predictive and were excluded from further evaluation, specifically: ‘AvgSpec’, ‘AvgP’, ‘MaxP’, ‘Fold_Change’, ‘BFDR’, ‘AvgSpec_nonciliated_bioid’, ‘AvgP_nonciliated_bioid’, ‘MaxP_nonciliated_bioid’, ‘Fold_Change_nonciliated_bioid’, ‘BFDR_nonciliated_bioid’.

An additional 6 features were deemed to carry no information (i.e. had insufficient training data to direct learning) and were ignored by the model: Hs_helaN_ph_hcw120_1_pq_euc’, ‘Hs_helaN_ph_hcw120_2_pq_euc’, ‘Hs_helaN_ph_saf48_pq_euc’, ‘Hs_helaN_ph_tcs375_P1_apex’, ‘Hs_helaN_ph_tcs375_P2_pq_euc’, ‘Hs_helaN_ph_tcs375_P3_pq_euc’. After cross validation, the top-ranked base classifier was the LightGBM model. AutoGluon’s ensemble model (i.e. WeightedEnsemble_L2) did not improve accuracy over the base LightGBM model. For a performance comparison of all AutoGluon models, see Supplemental Table 7.

The final model was used to predict confidence scores (i.e. DirectContact scores) of 25,992,007 unlabeled protein pairs for their probability of interacting directly (Supplemental Table 8). Based on the distribution of DirectContacts2 score (Supplemental Figure 2A), confidence thresholds of 0.7 and 0.9 were rationally selected to screen out lower confidence predictions. Thresholding at a DirectContacts2 score ≥ 0.7 yielded a set of 57,494 confident predictions, which we define to be our “full” DirectContacts2 network (Supplemental Table 9). Within this set, 24,328 pairs had a DirectContacts2 score ≥ 0.9 and were deemed “highly confident”.

### Precision Recall Analysis

Precision recall curves were calculated using the prcurve.py script in the protein_complex_maps package (https://github.com/KDrewLab/protein_complex_maps/blob/master/protein_complex_maps/eval uation/plots/prcurve.py [Commit: 3ebfb06]) using parameters --add_tiny_noise and --complete_benchmark. The script uses the Python sklearn package^48^ precision_recall_curve function and the matplotlib package^49^ for visualization. The model performance is evaluated on the leave-out set, consisting of 638 direct contacts (Supplemental Table 3) and 898 indirect contacts (Supplemental Table 4). We compared our new DirectContacts2 model as well as the original DirectContacts^17^ (https://doi.org/10.1371/journal.pcbi.1005625.s003), hu.MAP2.0^15^ (https://zenodo.org/records/15293715/files/humap2_ppis_ACC_20200821.pairsWprob.gz?down load=1), hu.MAP3.0^28^ (https://humap3.proteincomplexes.org/static/downloads/humap3/humap3_all_ppis_202305.pairs Wprob.gz), HuRI^26^ on a precision-recall basis and randomly shuffled predictions. HuRI was not utilized in the generation of the DirectContacts2 network and was used only for evaluation purposes. HuRI was downloaded from http://www.interactome-atlas.org/data/HuRI.tsv. HuRI interactions do not have an associated confidence score, so as done in hu.MAP2.0^15^ for precision recall analysis, HuRI interactions were ranked based on the number of separate assays in which the interaction was identified. We also evaluated our model using a modified set of negatives that included additional negatives from inter-complex indirect contacts (i.e. pairs of proteins in separate complexes) totaling 41,622 pairs. The inputs to the prcurve.py script were identical to above except for the --input_negatives parameter using the set of negatives from Supplemental Table 5, rather than from Supplemental Table 4.

### Comparison to AlphaFold2 modeled interactions

To evaluate DirectContact2’s ability to inform complex structural modeling, we compared our predictions to confidence values of AlphaFold2/AlphaFold2-Multimer-modeled structures downloaded from Burke et al.^14^ and Jänes et al^13^. AlphaFold2 models and confidence scores (i.e. pDockQ values) were downloaded from (https://archive.bioinfo.se/huintaf2/) and [https://ftp.ebi.ac.uk/pub/databases/ProtVar/predictions/interfaces/2024.05.28_interface_models _high_confidence.tar, https://ftp.ebi.ac.uk/pub/databases/ProtVar/predictions/interfaces/2024.05.28_interface_models _low_confidence.tar]. We recalculated pDockQ values for Jänes et al. models using the pDockQ.py script (https://gitlab.com/ElofssonLab/FoldDock/-/blob/main/src/pdockq.py) using the default interface distance cutoff of 8Å. This resulted in a final set of 277,807 AlphaFold2 and AlphaFold-multimer models with associated pDockQ scores. We again tested the DirectContacts2, DirectContacts, hu.MAP2.0, hu.MAP3.0, and HuRI networks on their ability to rank the set of pairs on their likelihood to interact. We utilized their pDockQ scores as an evaluative metric. Due to differences in each network’s confidence scores, we calculated the fraction of pairs with pDockQ scores ≥ 0.5 out of each network’s top-ranked K predictions, for K between 500 and 25,000 in increments of 500, generating a fraction-confident-at-K curve (Figure 4A). The same analysis was performed at a pDockQ score threshold of ≥ 0.23 (Supplemental Figure 2D).

### Generation of AlphaFold3 models

To assess DirectContacts2’s utility in guiding novel structure predictions in practice, we modeled 2,573 random pairs with a DirectContacts2 score ≥ 0.5 in AlphaFold3. 366 were generated using the AlphaFold Server (alphafoldserver.com)^7^, and 2,207 using the official public release of AlphaFold3 (github.com/google-deepmind/alphafold3) installed locally. AlphaFold3 predictions included only the two peptide chains comprising a given pair in their structures, with no post-translational modifications. AlphaFold Server runs were performed without manual seeding, equivalent to selecting a random integer as a seed. We used a model seed of 1 for structures predicted locally. For local predictions, the multiple sequence alignments and model inferences were performed separately utilizing AlphaFold3’s --norun_inference and --norun_data_pipeline flags to optimize resource usage across CPU and GPU environments. Aside from this, local AlphaFold3 runs were performed with default settings. Computation for local runs was performed on Indiana University’s Jetstream2 HPC cluster, for which resources were granted via the NSF Access program.

For each pairwise structure prediction, pDockQ was evaluated using the same methodology and parameters discussed above. We then used this set of pairs to produce the analysis shown in Figure 4B, where we show the fraction of pairs with pDockQ scores ≥ 0.5 at different thresholds of DirectContacts2 score.

### Inter-protein cross-links comparison analysis

To evaluate DirectContacts2 predictions with respect to protein crosslinking data, we compared DirectContacts2 predictions to inter-protein cross-links identified in Wheat *et al*.^34^ (Dataset_S02, sheet “Dataset S2A”, link: https://www.pnas.org/doi/suppl/10.1073/pnas.2023360118/suppl_file/pnas.2023360118.sd02.xlsx). Cross-linking mass spectrometry (XL-MS) captures covalently linked residues between protein chains in close spatial proximity, providing strong experimental evidence of direct physical contact within multi-protein assemblies. Multiple cross-links per protein pair were treated as a single cross-link identification between the protein pair, creating a non-redundant set with one crosslink per protein pair. Finally, to ensure cross-linked protein pairs are inter-protein links, identified peptides for each protein in the pair were compared to the other protein in the pair. If a match was found, this suggested a possible self-link and the cross-link pair was excluded from further analysis.

To evaluate the DirectContacts2 and hu.MAP3.0 networks on their agreement with cross-linked pairs, we compared each set to a dataset of XL-MS pairs taken from Wheat et al.^34^ For this analysis, we utilized the “full” DirectContacts2 network (Supplemental Table 9), and extracted the same number of top-ranking pairs from hu.MAP3.0 (i.e. the 57,494 top-ranking pairs), creating two equally sized sets of predictions. These sets were intersected with Wheat et al. to find their overlap with the XL-MS pairs. For both DirectContacts2 and hu.MAP3.0, we then sampled random pairs of proteins from the non-redundant cross-link dataset, which were then intersected with the sets of pairs from each network. This sampling and intersection was repeated 1,000 times to generate background distributions representing the tendencies to predict cross-linked protein pairs by random chance. Finally, we used the previously calculated intersection sets of both networks with the XL-MS set of pairs to estimate each network’s enrichment for cross-linked protein pairs against their random backgrounds, obtaining Z-scores representing each network’s enrichment for cross-linked pairs over random chance.

### Agreement of AlphaFold models and inter protein crosslinks

AlphaFold2 models of protein interactions HYOU1-HSPA5 and TMED5-TMED10 were used from Burke et al^14^. AlphaFold-multimer models of NEFL-NEFM and NEFM-VIM were created using the ‘colabfold_batch’ command line script developed as part of ColabFold^50^. For each protein pair, we downloaded FASTA sequences from Uniprot, constructed input files, and ran colabfold_batch with the following parameters: ‘--templates --num-recycle 3 --use-gpu-relax --model-type alphafold2_multimer_v3’. Colabfold_batch code was downloaded from the localcolabfold github repository (https://github.com/YoshitakaMo/localcolabfold). To generate higher stoichiometry models of NEFM-VIM interaction, we created an AlphaFold3^7^ model with default settings of expected overlapping segments (based on crosslinking data), specifically NEFM segments 85-176, 181-349, 350-498 and VIM segments 85-175, 176-338, 363-466 (https://zenodo.org/records/15634862/files/AF3_fold_nefm_vim_tails12_heads_2024_05_23_13_21.zip?download=1). This, therefore, restricts prediction to the expected dimer-of-dimer interface. We then aligned in PyMol^51^ two copies of the original 1:1 stoichiometry NEFM-VIM model onto the dimer-of-dimer interface model (https://zenodo.org/records/15634862/files/NEFM_VIM_higherorder_20240529.cif?download=1). Crosslinks were visualized using PyXlinkViewer^52^. PyXlinkViewer crosslink files are also available from the Zenodo repository.

### Calculation of pdockQ scores

To assess AlphaFold-multimer predictions, we calculated pDockQ score using the pdockq.py script downloaded from https://gitlab.com/ElofssonLab/FoldDock/-/blob/main/src/pdockq.py. The script takes the PDB output from AlphaFold-Multimer and uses the plDDT of the interface residues to evaluate the likelihood of interaction.

### Orofacial digital syndrome complex

Mutations for OFD1 were identified using the Uniprot variant viewer^53^. The orofacial digital syndrome (OFDS) complex was identified in hu.MAP2.0 (id: HuMAP2_00328, http://humap2.proteincomplexes.org/displayComplexes?complex_key=HuMAP2_00328). Pairs from the OFDS complex were ranked by DirectContact2 scores, where OFD1-FOPNL and FOPNL-KIAA0753 were the highest scoring pairs within the complex. We used ‘localcolabfold’ (i.e. AlphaFold-multimer) to predict three dimensional structural models for these high-scoring protein pairs. The pDockQ score was calculated to evaluate likelihood of interaction based on the AlphaFold-multimer structures. Pathogenic mutations of OFD1 were mapped on the AlphaFold structure in PyMOL^51^. The trimer structural model of OFD1-FOPNL-KIAA0753 was created by aligning the common subunit of the dimer models (i.e. FOPNL) in PyMOL.

## Code and Data Availability

All code can be found at our GitHub repository: https://github.com/KDrewLab/DirectContacts2_analysis.git. The full feature matrix used for training the DirectContacts2 model can be found here: https://huggingface.co/datasets/DrewLab/DirectContacts2/blob/main/full/humap3_full_feature_matrix_20220625.csv.gz. The Autogluon model can be found on HuggingFace: https://huggingface.co/DrewLab/DirectContacts2_AutoGluon. AlphaFold structural models can be found on Zenodo (https://doi.org/10.5281/zenodo.15634861) and are available in ModelArchive^54^ (modelarchive.org) with the accession code ‘ma-iicqo’ (https://www.modelarchive.org/doi/10.5452/ma-iicqo).

## Author Contributions

KD conceived of the study. ERC, MDWG, and KD designed experiments, analyzed data, wrote software, discussed and interpreted the results, and wrote the manuscript. SNF analyzed data, wrote software, discussed and interpreted results, and edited the manuscript.

## Acknowledgments

KD is supported by National Institute of Child Health and Human Development (R00 HD092613) and BBSRC - National Science Foundation/Directorate for Biological Sciences [BB/X002179/1][BB/X002683/1][NSF2314278]. Computational resources were provided by DOE Argonne ALCF Director’s Discretionary (DD) allocation award (Polaris HPC), the NSF ACCESS Exchange [BIO250091], and University of Indiana JetStream2. We would also like to thank Jill Paojin Pan and Eunseo Kim for help with generating AlphaFold3 models.

## Supplemental Figures

**Supplemental Figure 1:**
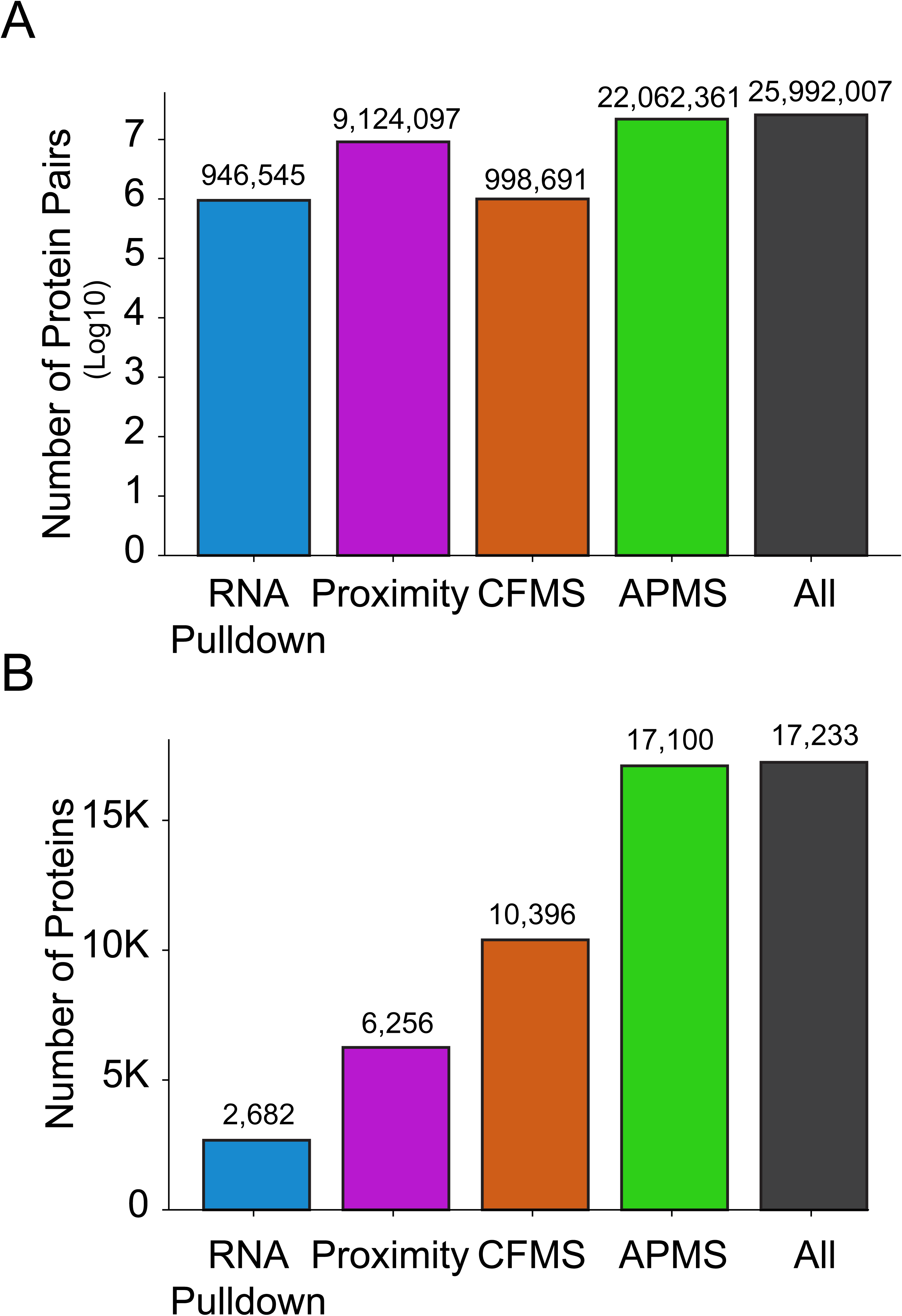
Proteomics dataset coverage of DirectContacts2. **A**. Coverage of each proteomic experimental evidence type for human protein pairs. **B.** Coverage of each proteomic experimental evidence type for human proteins.

**Supplemental Figure 2:**
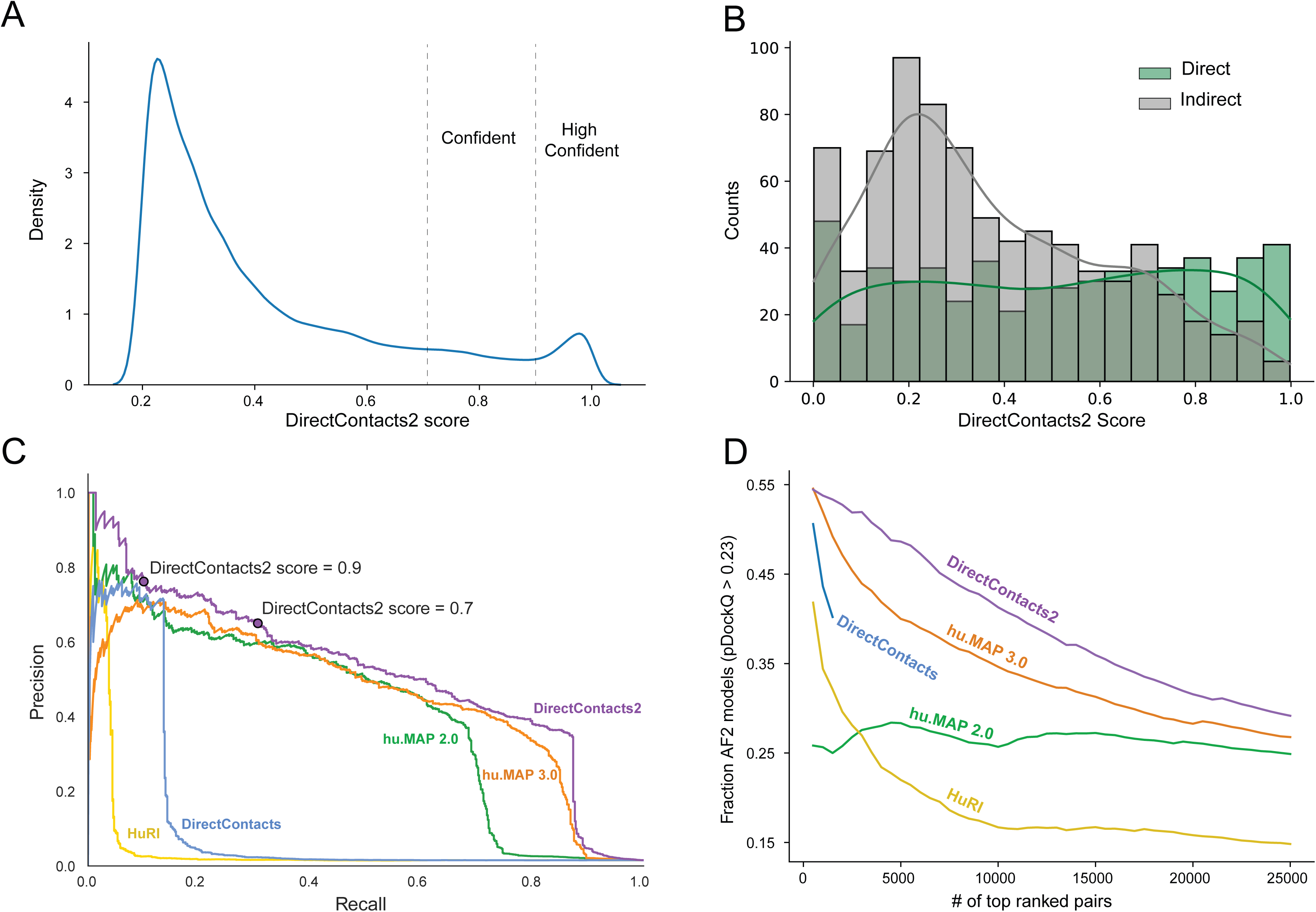
Evaluation of DirectContacts2 Network. **A.** Distribution of DirectContacts2 scores for all predicted pairs with score > 0.2. The distribution shows a bimodal distribution highlighting confident (>0.7) and highly confident (>0.9) predicted protein pairs. **B**. Distribution of DirectContacts2 scores for true direct interactions (green) and false indirect interactions (gray) in the leave-out test set derived from PDB structures of protein complexes. **C.** Precision-recall curve generated using the test set with inter-complex negative protein pairs (i.e. from separate complexes). DirectContacts2 score thresholds ≥ 0.7 (confident) and ≥ 0.9 (highly confident) are marked. **D**. Comparison of enrichment for high-confidence pair interfaces out of the set of top confident pairs for DirectContacts2 and other networks. For each set, the fraction of high-confidence interfaces (pDockQ ≥ 0.23) out of their top predictions was calculated for increasing numbers of top predictions.

**Supplemental Figure 3:**
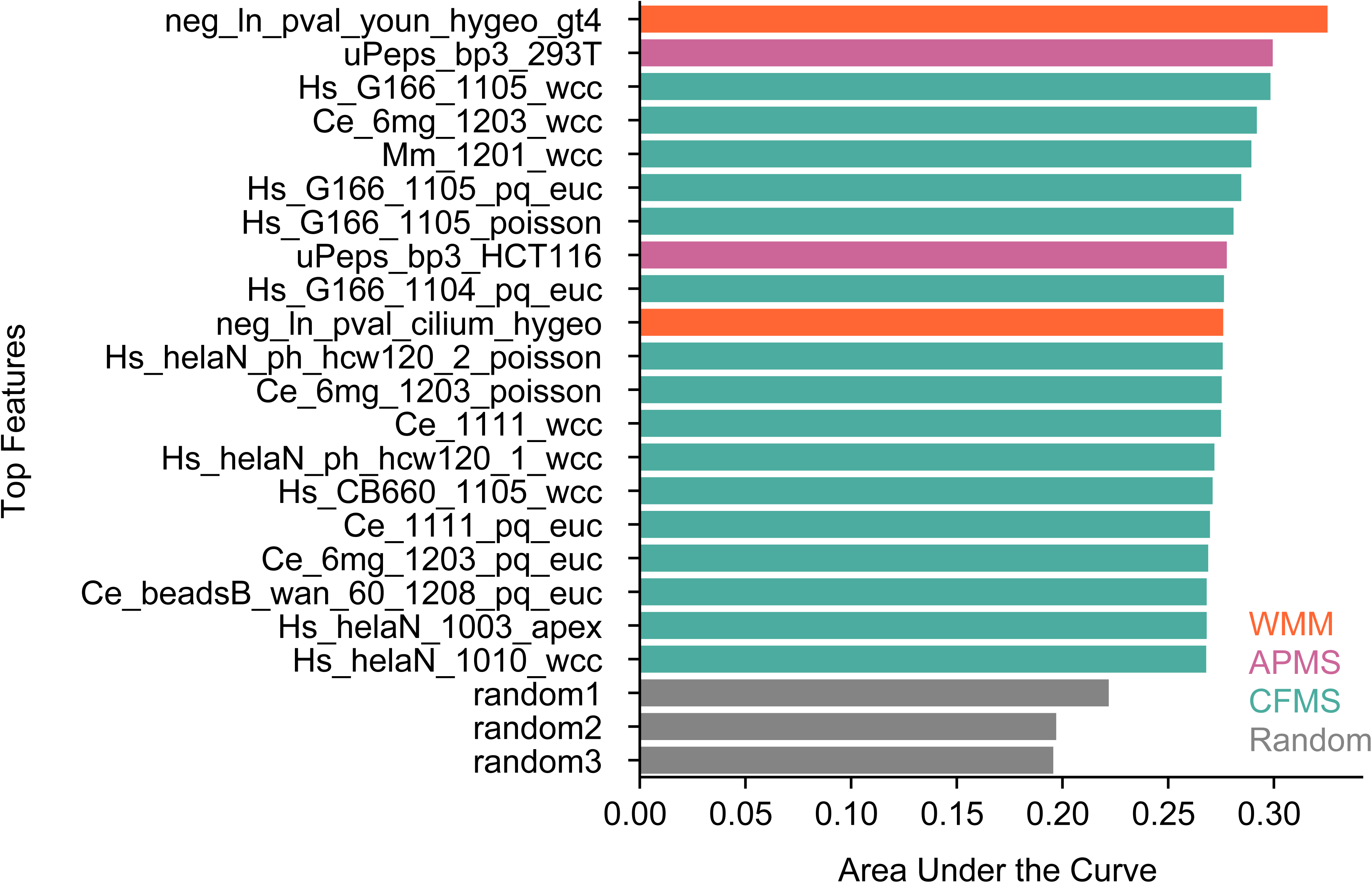
Area under the precision recall curve for top performing features to discriminate between direct and indirect interactions. Bar plot shows the area under the precision recall curve for untrained individual features evaluated on a training set excluding intercomplex negative edges. CFMS features are shown in green, APMS features are shown in pink, and WMM features are shown in orange. The two high ranking WMM features (neg_ln_pval_youn_hygeo_gt4 and neg_ln_pval_cilium_hygeo) are from proximity labeling experiments. Three random shufflings are also evaluated (gray).

**Supplemental Figure 4:**
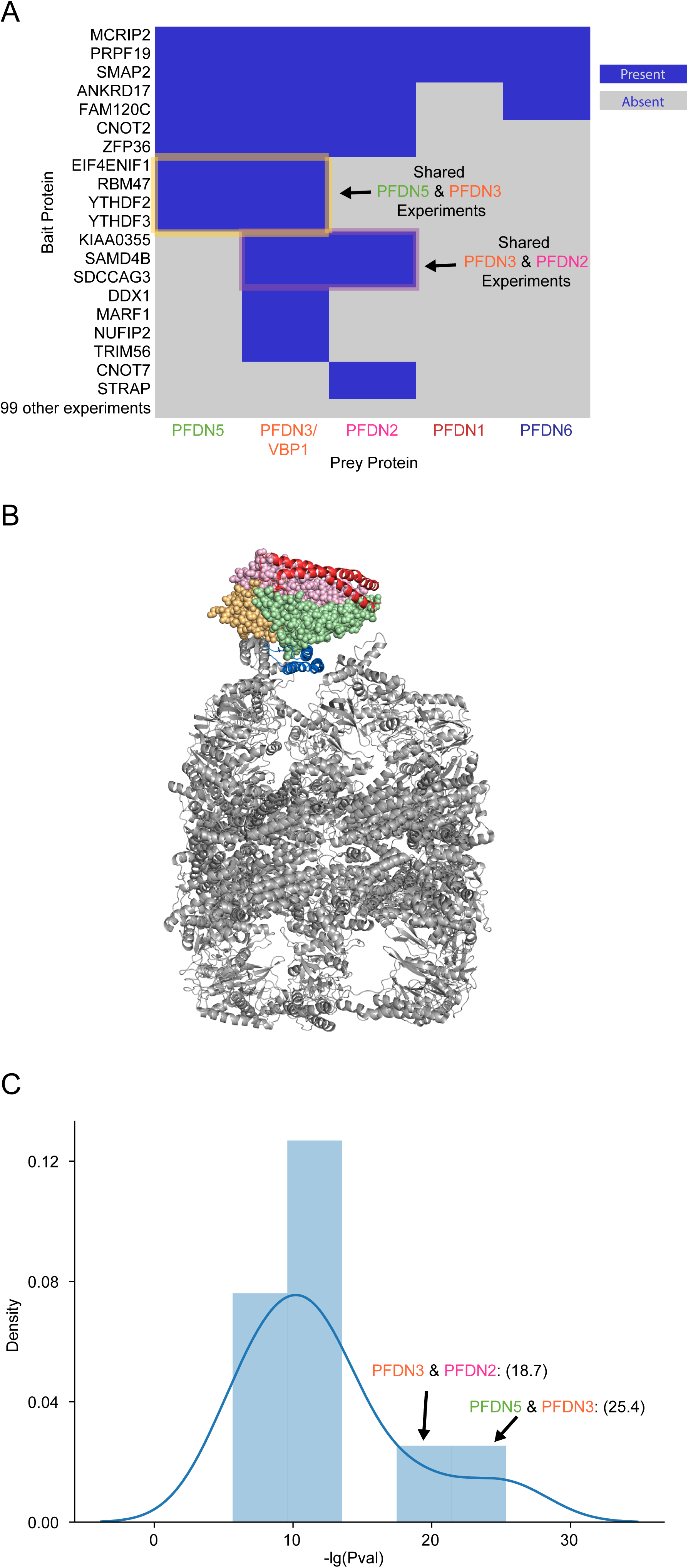
Weighted Matrix Model applied to proximity labeling experiments provides evidence for direct interactions in CCT/Prefoldin complex. **A.** Heatmap of proximity labeling experiments which observe presence of prefoldin subunits. Directly interacting pairs (PFDN5-PFDN3 (yellow box) and PFDN3-PFDN2 (purple box)) are seen together in the same experiments. **B.** PDB structure of CCT/Prefoldin complex showing PFDN5-PFDN3 and PFDN3-PFDN2 directly interact. CCT subunits in gray, PFDN5 (green), PFDN3 (orange), PFDN2 (pink), PFDN1 (red), PFDN6 (blue). **C.** Distribution of WMM calculated −lg(pval) of prefoldin subunit pairs shows bimodal distribution. Directly interacting pairs PFDN3-PFDN2 and PFDN5-PFDN3 are right shifted with higher WMM values.

## Supplemental Tables(csv files with headers or excel files)

Supplemental Table 1: Benchmark Training Positive Pairs

Supplemental Table 2: Benchmark Train Negative Pairs

Supplemental Table 3: Benchmark Test Positive Pairs

Supplemental Table 4: Benchmark Test Negative Pairs

Supplemental Table 5: Benchmark of Test Inter-Complex Negative Pairs

Supplemental Table 6: Benchmark Positive Pairs with PDBs

Supplemental Table 7: AutoGluon leaderboard of models

Supplemental Table 8: All 26 million predictions

Supplemental Table 9: Confident DirectContacts2 Predictions

Supplemental Table 10: AlphaFold3 model prediction metrics

